# Primary role of the Tol-Pal complex in bacterial outer membrane lipid homeostasis

**DOI:** 10.1101/2024.05.08.593160

**Authors:** Wee Boon Tan, Shu-Sin Chng

## Abstract

Gram-negative bacteria are defined by an outer membrane (OM) that contributes to envelope integrity and barrier function. Building this bilayer require proper assembly of lipopolysaccharides, proteins, and phospholipids, yet how the balance of these components is achieved is unclear. One system long known for ensuring OM stability is the Tol-Pal complex, which has been implicated in maintaining OM lipid homeostasis. However, assignment of Tol-Pal function has been challenging, owing to its septal localization and associated role(s) during division. Here, we uncouple the function of Tol-Pal in OM lipid homeostasis from its impact on cell division in *Escherichia coli*, by engineering a chimeric complex that loses septal enrichment. We demonstrate that this peripherally-localized Tol-Pal complex is fully capable of maintaining lipid balance in the OM, thus restoring OM integrity and barrier. Our work establishes the primary function of the Tol-Pal complex in OM lipid homeostasis, independent of its role during division.

## Introduction

The unique feature of the Gram-negative bacterial cell envelope is the presence of the outer membrane (OM). This asymmetric lipid bilayer is essential for growth and confers protection against external insults, including antibiotics. Lipopolysaccharides (LPS) are packed tightly together in the outer leaflet to impede permeability, while phospholipids (PLs) predominantly occupy the inner leaflet ^1,2^. The OM also contains abundant ꞵ-barrel proteins forming channels that allow selective passage of solutes across the bilayer ^3^. Notably, lipid-mediated interactions may facilitate the lateral clustering of OM proteins (OMPs) into islands that are extensively found throughout the OM ^4,5^. Consequently, the stability and barrier function of the OM depends acutely on the proper assembly of LPS, PLs, OMPs, and even lipoproteins, at optimal levels to achieve homeostasis. Furthermore, the OM is tethered to the peptidoglycan cell wall to maintain overall structural integrity and stability of the entire cell envelope ^6–8^.

The transport and assembly of LPS, OMPs, and lipoproteins into the OM are relatively well-characterized ^1,3,9^. These processes are known to be unidirectional, making it a challenge for the cell to directly adjust the levels of LPS, OMPs, and lipoproteins post assembly. In contrast, PL transport is bidirectional ^10–12^. This feature may facilitate the fine control of PL levels at the OM relative to other components, possibly allowing maintenance of a stable membrane ^13^. Unfortunately, the processes that move PLs between the two membranes are still poorly understood. The machines that mediate anterograde (IM-to-OM) transport of bulk PLs have remained elusive until recently, where three collectively essential AsmA family proteins, YhdP, TamB, and YdbH, are now implicated in the process ^14–16^. The OmpC-Mla system is known to transport PLs that have mislocalized to the outer leaflet of the OM back to the IM ^17^, yet whether it has any major impact on the overall OM stability is unclear. In fact, another system, the Tol-Pal complex, has also been implicated in retrograde (OM-to-IM) transport, but of bulk PLs ^13^. Unlike cells lacking OmpC-Mla, those lacking the Tol-Pal complex exhibit excess PL build-up at the OM, directly altering PL and LPS balance, and display slower measured rate of movement of OM PLs back to the IM ^13^. In this regard, the Tol-Pal complex may be a key player in maintaining OM integrity and stability via lipid homeostasis.

The Tol-Pal complex is highly conserved and comprises five proteins ^18–23^. TolQ, TolR, and TolA are integral membrane proteins that form a proton motive force (pmf)-utilizing subcomplex at the IM ^24^. Pal is an OM lipoprotein that can bind the periplasmic protein TolB or peptidoglycan (but not simultaneously) ^25,26^, with the latter interaction contributing to OM-cell wall tethering ^26–28^. Strains lacking any Tol-Pal component are known to have pleiotropic OM defects, including hypersensitivity to detergents and antibiotics, leakage of periplasmic contents, and OM vesiculation ^19,29–32^. These phenotypes indicate substantial perturbations to envelope integrity, which is consistent with lipid dyshomeostasis observed in the OM ^13,31,33^. The definitive assignment of Tol-Pal function is confounded, however, by the fact that Δ*tol-pal* mutants also exhibit division phenotypes, specifically cell chaining in extreme osmolality ^34,35^. It is believed that the lack of Tol-Pal causes delayed OM invagination ^34,36^, and most recently, also cell wall remodeling and separation defects ^37^. Notably, all components of the Tol-Pal complex localize to the cell septum during division ^34,36,38,39^. Septally localized TolQRA facilitates the recruitment of TolB and Pal to the division site ^36,38^, where pmf-induced conformational changes in TolA are thought to transmit a pulling force on TolB. Consequently, TolB-Pal interaction can be perturbed, leading to the dynamic enrichment of Pal-peptidoglycan binding, believed to be required for proper OM invagination at mid-cell. While TolQRA is also thought to regulate cell wall biosynthesis ^40^, it is not clear how enrichment of the Tol-Pal complex at the division site may ultimately modulate peptidoglycan remodeling, and impact cell separation. It is also unclear if, and how, septal localized Tol-Pal contributes to OM lipid homeostasis. Yet, the involvement of Tol-Pal in both pathways is evident from synthetic genetic interactions between the *tol-pal* locus and genes involved in either cell wall remodeling or OM biogenesis ^32^. Interestingly, alleviating cell wall separation and division defects in *tol-pal* mutants does not appear to rescue OM instability ^37^. Given the complex phenotypes, teasing out the exact function(s) of Tol-Pal in these two seemingly independent processes has been challenging.

One approach to uncouple the role of the Tol-Pal complex in OM lipid homeostasis from cell division would be to prevent septal localization. Since PLs occupy the entire inner leaflet of the OM, it is possible that the Tol-Pal complex remains functional all around the cell envelope, in its proposed role in retrograde PL transport ^13^. To test this idea, we explore the possibility of a chimera complex based on the known cross-compatibility between TolQR-TolA and the homologous ExbBD-TonB system. Like TolQRA, ExbBD uses the pmf to induce conformational changes in TonB to exert a mechanical pulling force, but on TonB-dependent transporters for metal/siderophore uptake ^41–44^. Both systems are also hijacked by bacteriocins for cell entry. Even though ExbBD-TonB does not display preference for the septal region ^45^, some degree of cross complementation has been reported between ExbBD and TolQR ^46–50^, indicative of possible interchangeability across the IM subcomplexes. In this work, we engineer a chimeric version of TonB^TM^-TolA that can work with the ExbBD subcomplex in *Escherichia coli*, giving rise to an assembled IM complex that no longer localizes specifically to the cell division site. We show that cells only expressing this peripherally localized complex still display defects in division. Importantly, the ExbBD-TonB^TM^ -TolA complex is fully functional in maintaining OM lipid homeostasis, in a manner dependent on TolB and Pal. Our work establishes that the primary function of the Tol-Pal complex is to mediate OM lipid homeostasis in Gram-negative bacteria, independent of septal localization.

## Results

### A TonB^TM^-TolA fusion retains the functionality of TolA in maintaining OM integrity but not cell division in a manner dependent on ExbBD

We hypothesized that Tol-Pal maintains OM integrity, presumably via retrograde PL transport, independent of its roles during cell division, and this function does not require septal localization. To test this idea, we sought to engineer a TonB^TM^-TolA chimera that works with the ExbBD complex instead of TolQR. Notably, the ExbBD-TonB complex is not recruited to the division site ^45^, therefore the ExbBD-TonB^TM^-TolA variant may lose septal localization yet retain TolA function (Figure 1A). Since TolA or TonB primarily interacts with TolQR or ExbBD, respectively, via its N-terminal transmembrane helix, we replaced the transmembrane helix of TolA with that of TonB. In this TonB^TM^-TolA chimera, the first 34 residues of TonB are fused to the periplasmic domains of TolA (35-421 a.a.) (Figure 1B). To assess its function in OM integrity, we expressed this construct in a MG1655 Δ*tolQRA* background strain, and examined SDS/EDTA or vancomycin sensitivity. Additional copies of ExbBD were expressed *in trans* in this mutant, as well as all control strains, to ensure sufficient ExbBD proteins available for both native TonB and TonB^TM^-TolA. As expected, the Δ*tolQRA* strain was extremely sensitive to SDS/EDTA and vancomycin. Remarkably, we found that the presence of the TonB^TM^-TolA fusion protein conferred strong resistance to both insults, indicating a fully restored OM barrier (Figure 2A, 2B). In contrast, wild-type TolA only marginally rescued the barrier defects when expressed at comparable levels (Figure S1). These observations are not unexpected, given that ExbBD is only known to weakly complement the absence of TolQR ^46,47,51^, and presumably less able to complex with TolA. Nevertheless, the striking difference between TonB^TM^-TolA and TolA highlights the importance of the TonB transmembrane helix (to effectively complex with ExbBD). Mutants lacking functional Tol-Pal also leak periplasmic contents, including RNase I, as indicated by the presence of RNA-degrading activity in culture supernatants (Figure 2C). Expression of the TonB^TM^-TolA construct fully prevented RNase I leakage (Figure 2C), further confirming that the chimera protein effectively replaces the role of TolA in maintaining OM integrity and barrier function.

**Figure 1.**
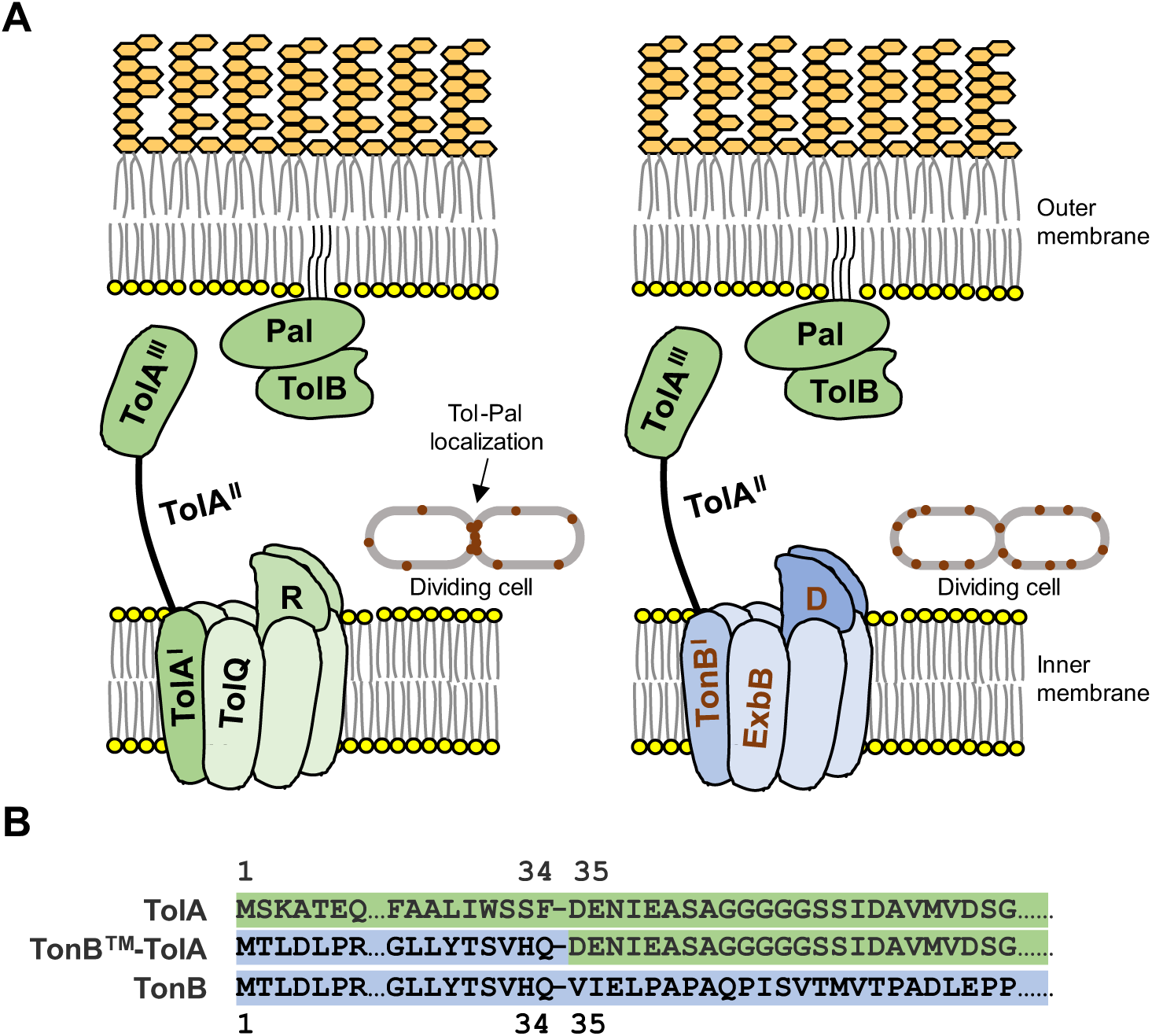
Preventing septal enrichment of the Tol-Pal complex using a TonB^TM^-TolA chimera. (A) Schematic representation of the native Tol-Pal complex (*left*) and the engineered chimeric complex (*right*), and their expected intracellular localization in dividing cells. Native Tol-Pal contains the TolQRA subcomplex at the IM, while the chimeric version contains instead a TonB^TM^-TolA fusion that works with ExbBD in a subcomplex that is not recruited to the division site. TolB and Pal remain the same in both scenarios. (B) Design of the TonB^TM^-TolA fusion protein. The TonB^TM^-TolA construct contains the first 34 amino acids from the N-terminus of TonB, including its transmembrane helix, fused directly to the periplasmic domains of TolA (amino acid 35-421).

**Figure 2.**
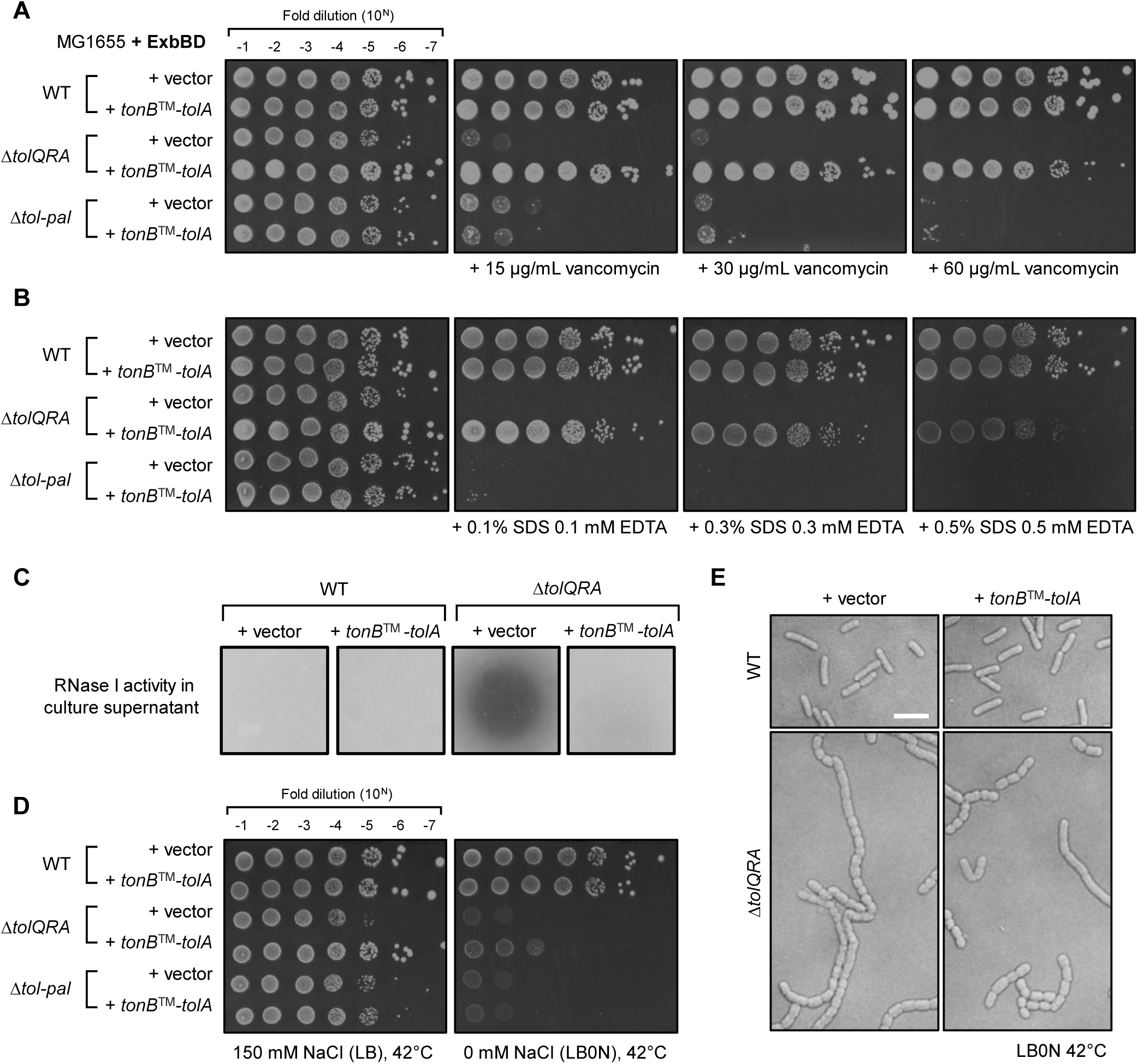
Chimeric ExbBD-TonB^TM^-TolA preserves the function of TolQRA for outer membrane integrity but not cell division. (A, B) Efficiency of plating (EOP) of MG1655 wild-type (WT), Δ*tolQRA*, and Δ*tol-pal* strains, containing either the pET23/42 empty vector ^68^ or the plasmid expressing TonB^TM^-TolA, on LB agar plates supplemented with (A) vancomycin or (B) SDS/EDTA at the indicated concentrations at 37°C. (C) RNase leakage assay of the WT and Δ*tolQRA* strains expressing TonB^TM^-TolA chimera, as judged by degradation of RNA (halo formation) when filtered culture supernatants were spotted on agar containing yeast RNA. (D) EOP of the same strains in (A) and (B) on either normal LB (150 mM NaCl) or LB0N (no NaCl) agar at 42°C. (E) Differential interference contrast (DIC) microscopy images of WT and Δ*tolQRA* strains expressing TonB-TolA chimera grown in LB0N media at 42°C. Scale bar represents 5 µm. All strains used here express additional copies of ExbBD from a pBAD33 vector ^69^ to enhance ExbBD-TonB^TM^-TolA chimera complex formation, and to avoid depletion of ExbBD for its native TonB-dependent function.

Since TolQ and TolR were absent in this strain, the recovery of Tol-Pal OM function must be derived from an ExbBD-TonB^TM^-TolA complex. Consistently, when the chromosomal copy of *exbD* was removed, TonB^TM^-TolA no longer confer SDS/EDTA and vancomycin resistance in Δ*tolQRA* (Figure S2). Furthermore, a mutation (H20A) within the transmembrane helix of TonB, known to affect ExbBD-TonB complex formation and pmf utilization ^52–54^, fully abolished the ability of TonB^TM^-TolA to maintain OM barrier function (Figure S2). To date, all reported functions of Tol-Pal require every component of the complex (TolQ, TolR, TolA, TolB, and Pal) to be present; thus, we checked if the ExbBD-TonB^TM^-TolA chimera complex require TolB and Pal using a Δ*tolQRA-tolB-pal* (Δ*tol-pal*) deletion strain. In this background further lacking TolB and Pal, TonB^TM^-TolA was unable to restore OM barrier function (Figure 2A, 2B), confirming that the fusion protein is performing a native Tol-Pal function.

Next, we checked if the TonB^TM^-TolA chimera also retains TolA function during cell division. As previously reported ^37^, the Δ*tolQRA* mutant was sensitive to low osmolarity media (LB0N) at elevated temperatures (Figure 2D), a phenotype associated with cell chaining division defects. Interestingly, expressing TonB^TM^-TolA did not rescue growth, nor fully prevent cell chaining defects, under this condition (Figure 2D, 2E). Therefore, the ExbBD-TonB^TM^-TolA complex appears incapable of replacing TolQRA in septal cell wall remodeling and/or OM invagination. Importantly, TonB^TM^-TolA was still stably expressed in cells grown at the same elevated temperature, thus effectively restoring OM barrier function (Figure S3). Overall, our findings demonstrate that the TonB^TM^-TolA fusion, in complex with ExbBD, is able to execute the native function of Tol-Pal in maintaining OM stability, but not in cell division processes.

### The ExbBD-TonB^TM^-TolA chimera complex ‘loses’ septal localization and ability to enrich Pal at mid-cell

The inability of TonB^TM^-TolA to rescue cell division defects may be due to lack of septal localization. To determine if this is true, we examined the cellular localization of an N-terminally sfGFP-tagged version of TonB^TM^-TolA. This sfGFP-TonB^TM^-TolA functioned similarly to the TonB^TM^-TolA construct (Figure S4). As a control, we also made a sfGFP-TolA fusion (with native TolA transmembrane domain), which did not rescue Δ*tolQRA* phenotypes (like wild-type TolA (Figure S1)), despite being expressed at similar levels compared to sfGFP-TonB^TM^-TolA (Figure S4). In both WT and Δ*tolQRA* strains, sfGFP-TolA was strongly enriched at cell division sites as expected. In stark contrast, sfGFP-TonB^TM^-TolA did not localize specifically to the cell septum but was randomly distributed around the cell periphery of both background strains (Figure 3A, 3B, 3C). Therefore, forcing TonB^TM^-TolA to interact with ExbBD prevents septal localization.

**Figure 3.**
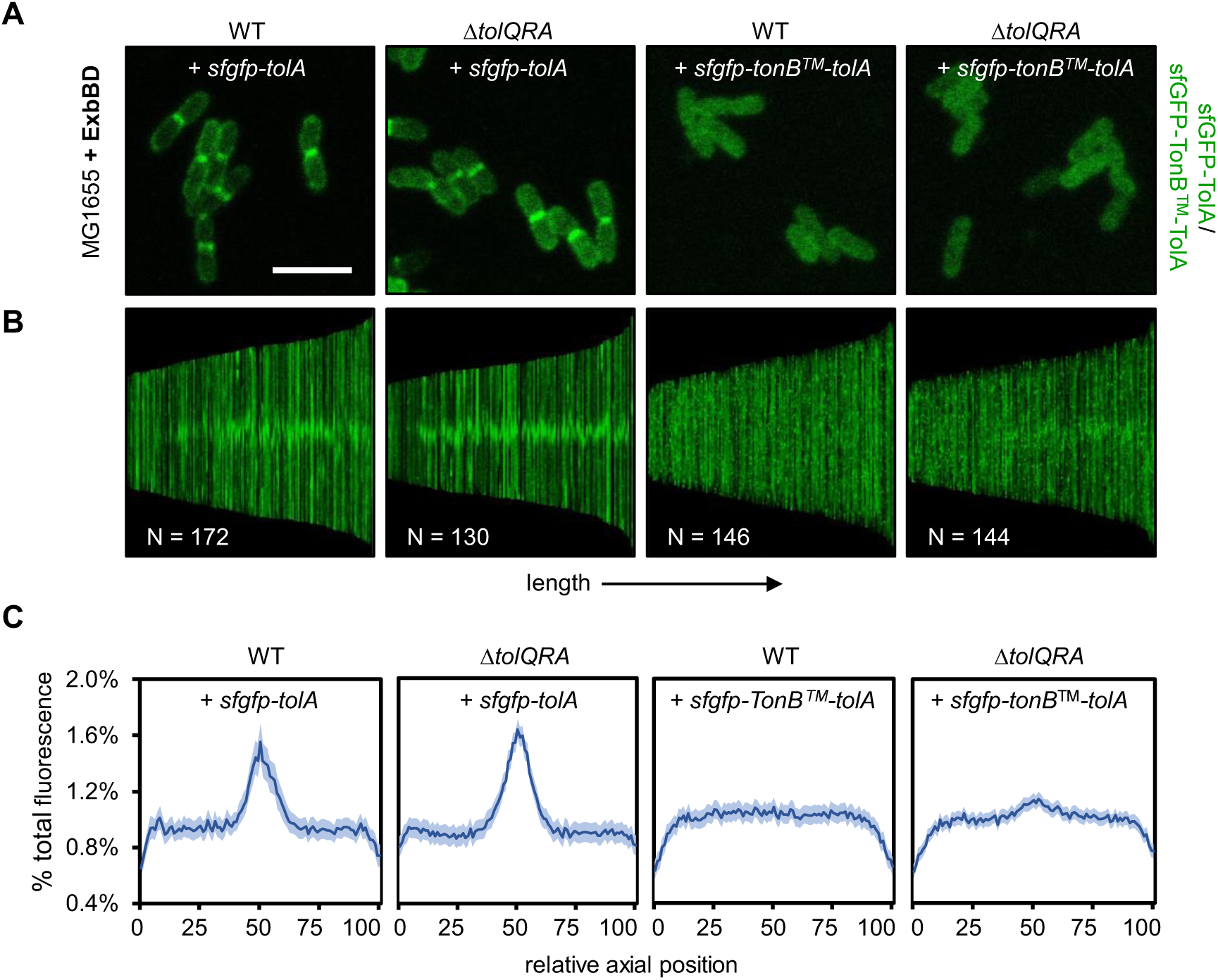
TonB^TM^-TolA is not spatially enriched at the cell septum. (A) Fluorescence microscopy images of MG1655 WT and Δ*tolQRA* strains, expressing sfGFP-TolA or sfGFP-TonB^TM^-TolA from pET23/42. Scale bar represents 5 µm. (B) Demographic representations of the sfGFP-TolA or sfGFP-TonB^TM^-TolA fluorescence signals in the indicated strains measured and normalized along the long axis of individual cells, sorted according to cell length and aligned to mid-cell. (C) Average percentage distribution of the sfGFP-TolA or sfGFP-TonB^TM^-TolA fluorescence signals along the long axis of all cells measured in (B), where the relative axial position of 50 corresponds to mid-cell (septum), and 1 and 100 correspond to the poles, respectively. 95% confidence intervals are shown. All strains here express additional copies of ExbBD from a pBAD33 vector.

TolQRA is believed to dislodge TolB for Pal enrichment at the septum ^36^, therefore we also investigated Pal localization in the presence of the ExbBD-TonB^TM^-TolA complex. Here, we expressed sfGFP-TonB^TM^-TolA and ExbBD in a previously reported Δ*tolQR* strain that also produces a functional Pal-mCherry fusion (from the native *tol*-*pal* locus) ^38^. In the WT background, Pal-mCherry was enriched at division sites due to the presence of functional TolQRA (Figure 4A, 4B, 4C). In the Δ*tolQR* background, accumulation of septal Pal-mCherry was largely abolished, as others have observed ^38^. Remarkably, we demonstrated that the presence of the ExbBD-TonB^TM^-TolA complex did not restore septal Pal enrichment (Figure 4); we conclude that the observed cell division defects (Figure 2D, 2E, S2, S4) in strains expressing ExbBD-TonB^TM^-TolA were due to loss of septal-localized Tol-Pal function. Given that OM barrier functions are fully restored, we believe that the ExbBD-TonB^TM^-TolA complexes localized peripherally are sufficient to perform the function of Tol-Pal in OM stability and homeostasis.

**Figure 4.**
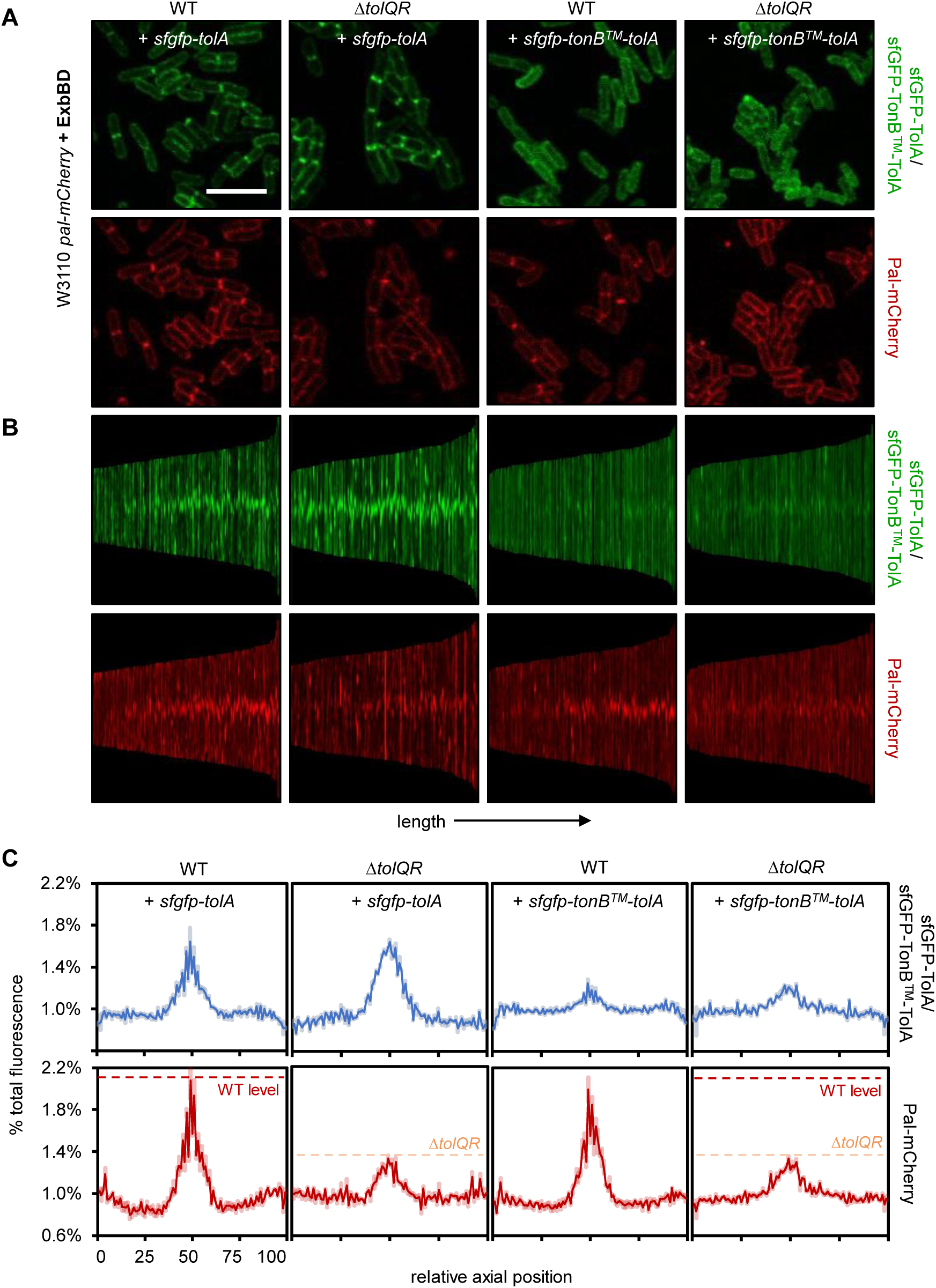
Peripherally localized ExbBD-TonB^TM^-TolA complex no longer promotes septal enrichment of Pal. (A) Fluorescence microscopy images of W3110 WT (expressing Pal-mCherry from the native *pal* chromosomal locus) and the corresponding Δ*tolQR* strains, expressing sfGFP-TolA (control) or sfGFP-TonB^TM^-TolA. Scale bar represents 5 µm. (B) Demographic representations of the sfGFP-TolA/sfGFP-TonB^TM^-TolA (*green*) and Pal-mCherry (*red*) fluorescence signals in the indicated strains measured and normalized along the long axis of individual cells, sorted according to cell length and aligned to mid-cell. (C) Average percentage distribution of the sfGFP-TolA/sfGFP-TonB-TolA (*blue*) and Pal-mCherry (*scarlet*) fluorescence signals along the long axis of all cells measured in (B), where the relative axial position of 50 corresponds to mid-cell (septum), and 1 and 100 correspond to the poles, respectively. 95% confidence intervals are shown. All strains here express additional copies of ExbBD from a pBAD33 vector.

An unexpected observation in our microscopy studies was that expressing ExbBD-TonB^TM^-TolA actually rescued the mild morphological defects seen in the above Δ*tolQRA* and Δ*tolQR* strains grown under normal osmolality conditions (Figure 5, S5). Disruption of Tol-Pal function is known to cause moderate increase in cell width, giving rise to “fatter” cells ^31,34^, a morphological change previously attributed to defects in cell division. Since restoring OM integrity alone in these strains returned cell width back to normal, it is likely that this cell morphological defect arises from OM lipid dyshomeostasis in *tol-pal* mutants.

**Figure 5.**
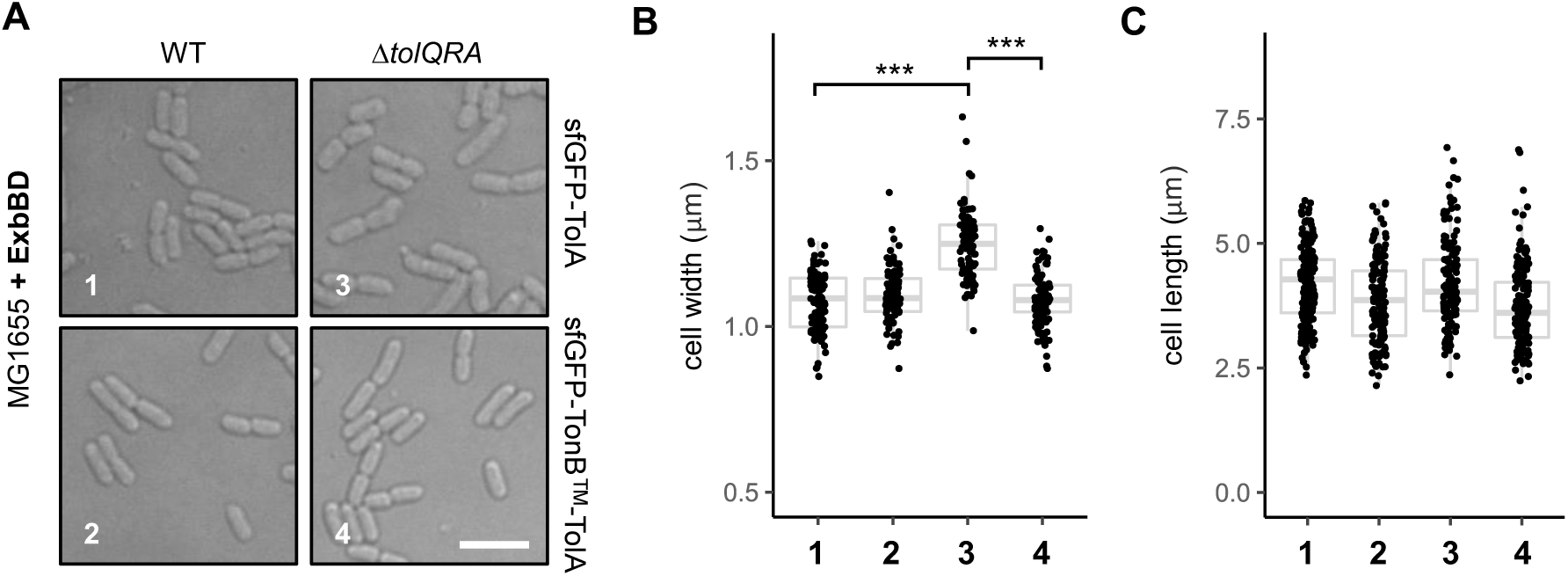
Cell morphological defects are restored by ExbBD-TonB^TM^-TolA. (A) DIC microscopy images of MG1655 WT and Δ*tolQRA* strains, expressing sfGFP-TolA or sfGFP-TonB^TM^-TolA. Scale bar represents 5 µm. (B, C) Quantification of the cell (B) width and (C) length of >100 individual cells of the same strains (*numbered*) shown in (A). Wilcoxon ranked sum test: ***, *P* < 0.0001. All strains here express additional copies of ExbBD from a pBAD33 vector.

### Peripherally localized ExbBD-TonB^TM^-TolA complex retains Tol-Pal function in OM lipid homeostasis

*tol-pal* mutants are known to release excess amounts of OM vesicles (OMVs) relative to WT ^19,31^, especially from the septum; this hypervesiculation phenotype is believed to be associated with both the loss of septal peptidoglycan-OM tethering, as well as OM instability and lipid dyshomeostasis. Since ExbBD-TonB^TM^-TolA rescues OM defects, we wondered if OM hypervesiculation was alleviated. Following [^14^C]-acetate labeling of cellular lipids, we isolated and quantified OMVs from culture supernatants of WT and Δ*tolQRA* strains expressing ExbBD and TonB^TM^-TolA. Interestingly, while Δ*tolQRA* mutant produced ∼50-fold more OMVs than WT, the presence of ExbBD-TonB^TM^-TolA reduced that excess by ∼79% (i.e. only ∼10-fold more than WT) (Figure 6A). This residual OM vesiculation is likely due to the lack of Tol-Pal function at the division site, manifesting as problems in OM invagination. Importantly, it appears that peripherally localized Tol-Pal function is sufficient to reduce OM hypervesiculation significantly, highlighting OM instability and lipid dyshomeostasis as the main contributors to this phenotype in *tol-pal* mutants.

**Figure 6.**
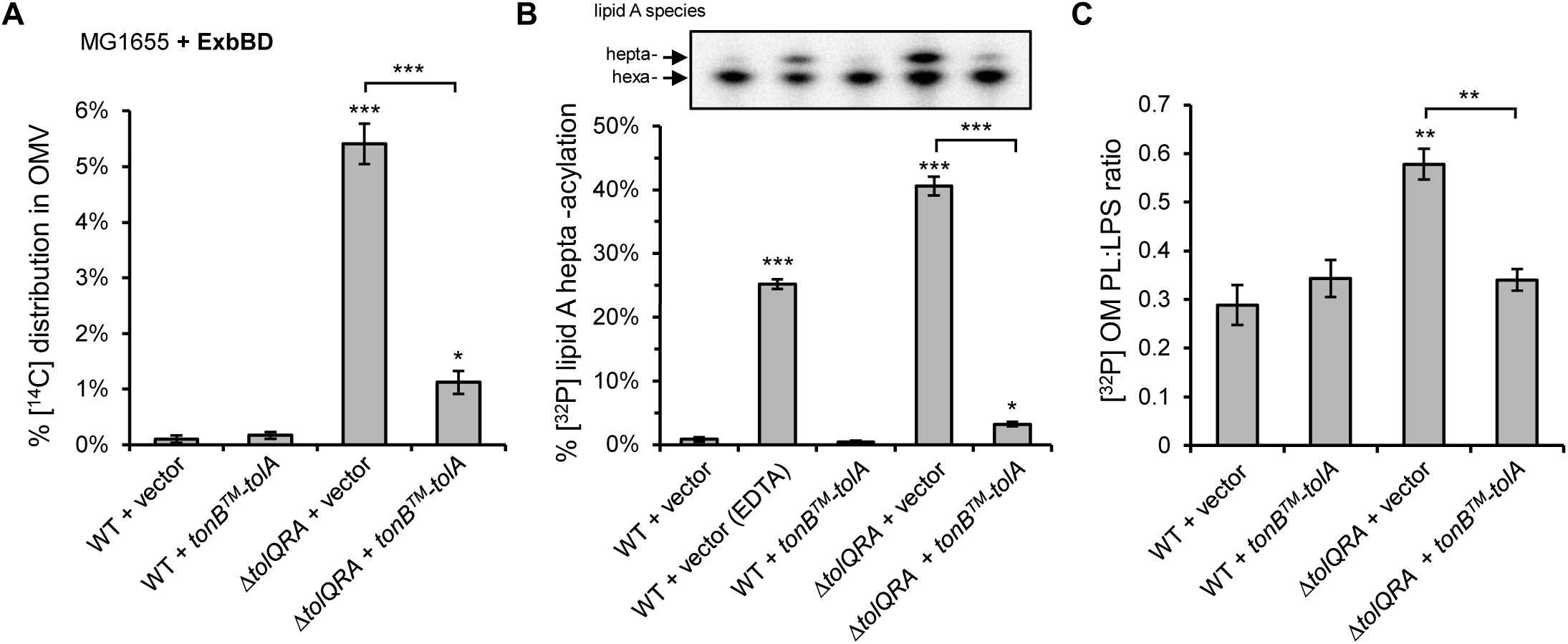
Peripherally localized Tol-Pal maintains outer membrane lipid homeostasis. (A) Quantification of OMV levels in MG1655 WT and Δ*tolQRA* strains expressing TonB^TM^-TolA, represented by levels of [^14^C]-labeled lipids in the filtered culture supernatant, relative to total [^14^C]-acetate labeling (cell pellet + OMV) of the respective strains. (B) Quantification of OM lipid asymmetry defects in WT and Δ*tolQRA* strains expressing TonB-TolA, represented by the levels of [^32^P]-labeled hepta-acylated lipid A, following thin-layer chromatography (inset)/phosphor imaging analyses. PagP-mediated transfer of an acyl tail from outer leaflet PLs to hexa-acylated lipid A serves as a proxy for lipid asymmetry in the OM. (C) Quantification of [^32^P]-labeled PL to LPS ratios in the OMs of WT and Δ*tolQRA* strains expressing TonB^TM^-TolA. Error bars represent standard deviations across three independent replicate experiments. Student’s *t*-tests: *, *P* < 0.01; **, *P* < 0.001; ***, *P* < 0.0001 (as compared to WT with empty vector, unless otherwise specified). All strains here express additional copies of ExbBD from a pBAD33 vector.

We went on to ascertain if ExbBD-TonB^TM^-TolA indeed restored OM lipid homeostasis, thereby rescuing OM barrier defects and hypervesiculation. Mutants lacking the Tol-Pal complex accumulate excess PLs in the OM, reflected in an elevated PL:LPS ratio ^13^. This PL excess presumably gives rise to perturbation in OM lipid asymmetry, where PLs mislocalize to the outer leaflet, providing substrates for hepta-acylation of LPS by PagP ^13^. Expectedly, in [^32^P]-radiolabeled lipid profiling experiments, the Δ*tolQRA* mutant exhibited increased PL:LPS ratio and hepta-acylated LPS levels, as previously reported (Figure 6B, 6C). Remarkably, we demonstrated that expression of TonB^TM^-TolA reversed both molecular phenotypes back to levels similar to WT, effectively re-establishing OM lipid homeostasis. Taken together, we conclude that the Tol-Pal complex can mediate OM lipid homeostasis without enrichment to the division site. Maintaining OM stability and homeostasis is the primary function of the Tol-Pal complex.

## Discussion

The Tol-Pal complex is functionally involved in cell division and in OM lipid homeostasis. In this work, we have demonstrated that the Tol-Pal complex performs its function in OM homeostasis independent of its roles during cell division. We have engineered a chimeric protein, with the transmembrane domain of TonB fused to the periplasmic “effector” domains of TolA, that forms a functional complex with ExbBD instead of TolQR. This ExbBD-TonB^TM^-TolA complex itself does not localize, hence cannot enrich Pal, to the septum (Figure 3, 4), resulting in the loss of cell division function(s) associated with the Tol-Pal complex (Figure 2D, 2E). Yet, this peripherally distributed chimera complex retains the function of Tol-Pal in maintaining OM lipid homeostasis (Figure 6), thus restoring OM stability and permeability barrier function (Figure 2A, 2B, 2C). Our work establishes that OM defects in *tol-pal* mutants are not indirect consequences of problem(s) that arises during cell division, and favors a model where the Tol-Pal complex plays a primary role in maintaining OM lipid homeostasis throughout the cell independent of septal localization.

The physiological function of the Tol-Pal complex has been a longstanding question in the field. As early as the 1960s, the Tol-Pal complex was thought to be important for OM stability ^18–20,30,55,56^. Strains lacking any of the Tol-Pal proteins release more OMVs and exhibit exacerbated OM permeability defects. Yet, it was also known that *tol-pal* mutants display cell chaining defects, but only under extreme osmolality conditions ^34,35^. Therefore, it has been challenging to interpret such pleiotropic phenotypes and assign the real function of the Tol-Pal complex. In 2007, the Tol-Pal complex was reported to localize to the cell septum ^34^, highlighting a potential role during cell division, specifically to facilitate OM invagination. This idea quickly gained traction then ^36,40,57,58^, given that Tol-Pal is a trans-envelope complex, and that Pal tethers the OM to the peptidoglycan layer; there appears to be an underlying assumption that the OM and cell chaining phenotypes in *tol-pal* mutants were somehow due to defective OM constriction during cell division. A key development that challenges this prevailing thought was the recent discovery of OM lipid dyshomeostasis in *tol-pal* strains ^13,33^, which can logically account for OM stability and permeability defects; it became less clear how defects in OM invagination can actually give rise to changes in lipid compositions in the OM. It is also now established that the cell chaining phenotype under low osmolality conditions is in fact due to defects in septal cell wall remodeling and separation ^37^. Notably, suppressing chaining phenotypes by overexpressing cell wall hydrolases did not significantly alleviate OM permeability and hypervesiculation defects in *tol-pal* strains ^37^. This is in agreement with OM phenotypes already manifesting in *tol-pal* strains grown under normal osmolality conditions where chaining is naturally suppressed. While it is unlikely that OM phenotypes are downstream of cell division defects in *tol-pal* mutants, it remained difficult to clarify true causal relationships. In this regard, we have now introduced a novel approach to uncouple the function of Tol-Pal in maintaining OM lipid homeostasis from its role(s) in cell division. By essentially preventing septal enrichment of a functional Tol-Pal complex, our strategy facilitates the definition of Tol-Pal function in OM lipid homeostasis, contributing to a critical advance in our understanding of this decades-old problem.

By separating the functions of the Tol-Pal complex in OM lipid homeostasis and cell division, we are also now able to explain most of the pleiotropic phenotypes observed in *tol-pal* mutants. OM permeability defects are a direct result of lipid dyshomeostasis. Interestingly, we have found that restoring OM lipid homeostasis was sufficient to rescue the mild morphological defects, i.e. wider cells, observed in *tol-pal* strains grown under normal osmolality conditions (Figure 5). Therefore, we reason that the wider cellular morphology is likely due to both decreased rigidity, and the increased surface area, of the OM, as a result of excess PL accumulation. Furthermore, it appears that OM hypervesiculation can also be largely attributed to lipid dyshomeostasis. We have shown that expressing the functional peripheral Tol-Pal complex was able to reduce OMV shedding drastically (Figure 6A), indicating that an unstable OM is indeed a major contributor to vesiculation in these cells. Overall, we deduce that defects in OM stability and permeability, hypervesiculation, and cellular morphology, in cells grown under normal osmolality, can be traced back to the primary role of the Tol-Pal complex in OM lipid homeostasis. Nonetheless, the Tol-Pal complex is still important for OM invagination during cell division. This conclusion is consistent with the residual OMV shedding present in cells with restored OM lipid homeostasis, but deficient in septal localization of Tol-Pal. However, it remains unclear if, and how, OM invagination defects may contribute to cell wall remodeling and chaining defects, observed only under extreme osmolality conditions. Given that changes in osmolality can exert additional stresses on the cell envelope, and notably also alter enzymatic activities of cell wall biosynthetic enzymes ^59^, remodeling and chaining defects may be a manifestation of ensuing synthetic interactions within the stressed envelope.

How the Tol-Pal complex mediates OM stability throughout the cell is not clear. We have demonstrated that retaining Tol-Pal function in the cell periphery is sufficient to maintain OM lipid homeostasis, but the molecular mechanism(s) remains elusive. Given the known ability of Pal to bind peptidoglycan, it is natural to first consider Pal-mediated OM-cell wall tethering as the main activity of the Tol-Pal complex. However, other tethering proteins (e.g. Lpp, OmpA) exist at much higher abundance throughout the cell in *E. coli* ^6,8,60^, raising the question how the lack of Pal-mediated tethering from the cell periphery would give rise to such severe OM defects observed in *tol-pal* mutants. Furthermore, removing only TolB, which is expected to increase Pal binding to cell wall, also does not alleviate the OM phenotypes. Alternatively, it may be that the more dynamic nature of this system contributes to OM stability, even when Tol-Pal function is peripherally distributed. TolQRA is believed to modulate the interaction between TolB and Pal, continually influencing the levels of Pal binding to cell wall, particularly at the cell septum ^36^. Yet, diffusion kinetics of Pal in the cell periphery of non-dividing cells did not change appreciably in response to TolA or TolB mutations that cause OM defects ^36^, suggesting little correlation between Pal-cell wall binding dynamics and Tol-Pal function. Overall, we believe that OM-cell wall tethering mediated by Pal is unlikely the key activity to realize Tol-Pal function in OM stability and lipid homeostasis. Instead, on the bases of excess PL accumulation in the OM and slower intermembrane PL exchange, we have previously hypothesized that the Tol-Pal complex has a possible activity in retrograde transport of bulk PLs ^13^. Since TolQRA harnesses the pmf for force generation across the cell envelope, in part to alter TolB-Pal interaction, speculative models for how this force can translate to PL transport activity have thus been proposed ^2,13^; these models include active shuttling of a yet-to-be identified PL-binding protein, and even pulling of the IM and the OM together for potential hemifusion events. In these scenarios, it is conceivable that Pal binding to peptidoglycan, which occurs through *meso*-diaminopimelate (*m*DAP) recognition ^27^, may be a way to locate non-crosslinked regions in the cell wall mesh for transenvelope transport. Ultimately, the proposed activity of the Tol-Pal complex in retrograde PL transport will enable fine control of PL content relative to other OM components, thereby directly contributing to the overall stability and homeostasis of the OM. These ideas require further investigation.

Regardless of the molecular mechanism, however, the function of the Tol-Pal complex in OM lipid homeostasis is likely also beneficial for OM invagination during cell division. After all, the IM components TolQ, TolR, and TolA are recruited to the septum via at least two independent ways linked to septal cell wall biosynthesis ^34,39,61^, indicating a possible evolutionary advantage for Tol-Pal function(s) to exist at mid-cell. While Pal-mediated tethering of the OM to the cell wall should contribute to septal OM invagination, we propose that changing lipid composition, i.e. membrane remodeling, would also be important for this process. Specifically, changes in PL content at the inner leaflet of the OM, possibly facilitated by retrograde PL transport, may modulate membrane curvature locally, in a way that ultimately leads to proper invagination. In this regard, it is likely that other lipid transport processes may also play unbeknownst role(s) to control septal membrane composition/curvature – an under-appreciated aspect of OM constriction during cell division. Future efforts should focus on elucidating the mechanisms of Tol-Pal function in OM lipid homeostasis and invagination.

## Materials and Methods

### Strains, plasmids, and growth conditions

List of *E. coli* strains used in this study can be found in the Supplementary Information. MG1655 strain expressing additional copies of ExbBD from pBAD33 vector was used as the wild-type strain for most experiments unless otherwise specified. W3110 strains expressing chromosomal Pal-mCherry from its native locus have been used in previous publications ^38^ and were acquired from the Duche laboratory. Gene deletion alleles were constructed using λ Red recombineering and transduced into relevant genetic background strains via P1 transduction ^62,63^. Strains were grown in Luria-Bertani (LB) broth (1% tryptone and 0.5% yeast extract, with or without 1% NaCl) supplemented with chloramphenicol (30 µg/ml), ampicillin (100 µg/ml), and/or 0.2% arabinose, to maintain plasmids or induce expression of ExbBD as needed. Agar plates contain 1.5% agar in the corresponding media. Cultures and plates were incubated at 37°C unless otherwise specified. Plasmids expressing ExbBD, TonB^TM^-TolA, sfGFP-TonB^TM^-TolA, sfGFP-TolA, and sfGFP were constructed using Gibson assembly ^64^ (detailed in Supplemental Information).

### Efficiency of plating

Sensitivity towards SDS/EDTA, vancomycin, and low salt were tested by spotting serially diluted cultures of the respective strains onto agar plates with the different conditions. When needed, agar plates were supplemented with chloramphenicol (30 µg/ml) and ampicillin (100 µg/ml) to maintain the plasmids and 0.2% arabinose to induce expression of ExbBD. Overnight cultures were 10-fold serially diluted in 150 mM NaCl on 96-well plate. Two (for SDS/EDTA plates) or five µl of each dilution were spotted onto the plates and incubated overnight at the indicated temperatures. All results shown are representative of at least three independent replicates.

### Periplasmic RNase leakage assay

Overnight cultures of the indicated strains were pelleted, and the supernatant filtered through 0.22 µM syringe filter. 10 µl of filtered supernatant were spotted onto agar plate containing spectinomycin (of which the strains were sensitive to) and 2 mg/ml yeast RNA. RNase leakages were visualized by overlaying cold 12.5% trichloroacetic acid onto the agar plates after overnight incubation at 37°C. Clear zone indicates lack of RNA and therefore the presence of RNase leakage into culture supernatant.

### Microscopy

To examine cellular morphology and fluorescence localization, overnight cultures were first diluted 1:2000 (1:1000 for LB0N experiment) into fresh media. The sub-cultures were allowed to grow to mid-log (∼4 hours) before being concentrated 10-fold by centrifuging at 3000 × *g* for 5 minutes. 5 µl of the concentrated cultures were spotted onto freshly made 1% agarose pad with the corresponding media. Images were acquired using a Zeiss LSM710 confocal microscope at 100x magnification and analysed using ImageJ (FIJI) and the MicrobeJ plugin ^65,66^.

### Quantification of OMV

Experiment to estimate OMV production had been previously described ^31^. Briefly, 10 ml cultures were grown in the presence of [1-^14^C]-acetate (final concentration of 0.2 µCi ml^-1^; Perkin Elmer NEC084A001MC) until OD_600_ reached ∼0.5-0.7, after which the culture supernatants were collected and filtered through 0.45 μm filters. The filtrates were ultracentrifuged, and the washed pellets were measured for [^14^C] radioactive counts on a scintillation counter (MicroBeta2®, Perkin-Elmer).

### PagP-mediated LPS hepta-acylation analysis

Experiment to quantify lipid asymmetry defects via measuring PagP-mediated lipid A palmitoylation (hepta-acylation) had been previously described ^13,67^. Briefly, 5 ml cultures were grown in the presence of [^32^P]-phosphate (final concentration of 1 µCi ml^-1^; Perkin Elmer NEX060001MC) overnight, after which the cell pellets were harvested and washed twice with Tris-buffered saline (TBS). The pellet was resuspended in single phase Bligh/Dyer mixture, incubated for 20 minutes at room temperature, and centrifuged at 21000 × *g* for 30 minutes. The pellets containing LPS were washed once with 1 ml of single-phase Bligh/Dyer mixture and subsequently resuspended in 0.45 ml 12.5 mM sodium acetate containing 1% SDS (pH 4.5). The resuspension was sonicated for 15 minutes and heated at 100°C for 40 minutes. The mixtures were then converted into two-phase Bligh/Dyer mixture and the lower phase (containing lipid A from hydrolyzed LPS) were collected and dried. The dried lipid A samples were resuspended in 30 μl of chloroform/methanol mixture (4:1) for thin-layer chromatography (TLC) analysis using Silica Gel 60 F254 TLC plate (Merck Millipore) with the solvent system consisting of chloroform/pyridine/96% formic acid/water (50/50/14.6/4.6). Dried TLC plate was exposed to phosphor storage screens and visualized. Spots were quantified and averaged based on three independent experiments of lipid A isolation.

### OM lipid composition analysis

Experiment to measure PL:LPS ratio at the OM had been previously described ^13,31,32^. Briefly, 10 ml cultures were grown in the presence of [^32^P]-phosphate (final concentration of 1 µCi ml^-1^; Perkin Elmer NEX060001MC) for until OD_600_ reached ∼1.5-2, after which the cell pellets were harvested and washed twice with Tris-buffered saline (TBS). Resulting cell pellets were re-suspended in 5 ml of 20% sucrose in 10mM Tris-HCl pH 8.0 (w/w) containing 1 mM PMSF and 50 μg ml^−1^ DNase I), and lysed by a single passage through a high-pressure French press (French Press G-M, Glen Mills) homogenizer at 8000 psi. Unbroken cells were removed by centrifugation at 3260 × *g* for 10 min. The cell lysates were subjected to a two-step sucrose gradient ultracentrifugation to separate to IM and OM fractions as previously described. OM fractions were pooled and concentrated for subsequent separation of PL and LPS. Protocol for the extraction of PLs and LPS had been described previously. Dried PLs were resuspended in 50 µl of a mixture of chloroform:methanol (2:1); while dried LPS pellets were resuspended in 50 µl 1% SDS. [^32^P] radioactivity of equal volumes of PL or LPS solutions were measured using scintillation counting (MicroBeta2®, Perkin-Elmer). Scintillation counts of OM PLs were divided by the counts of OM LPS to obtain the PL/LPS ratio. Data shown is representative of three independent triplicate experiments.

## Acknowledgments

We thank Denis Duché (CNRS, Aix-Marseille Université) for providing the W3110 *pal*-*mCherry* strains. This work was supported by the Singapore Ministry of Health National Medical Research Council under its Open Fund Individual Research Grant (MOH-000145) and the Singapore Ministry of Education Academic Research Fund Tier 1 grant (National University of Singapore-Faculty of Science Preparatory Grant Scheme) (both to S.-S.C.).

