## Supporting Information for "Primary role of the Tol-Pal complex in bacterial outer membrane lipid homeostasis"

*To whom correspondence should be addressed.

**ORCID ID:**

**This PDF file includes:**

Supplemental text
Figures S1 to S5

### Supplemental Information Text

#### Strain list

| Strain | Reference |
|---|---|
| MG1655 | - |
| MG1655 Δ*tolQRA*::*kan*$^{R}$ | Tan & Chng (2022) |
| MG1655 Δ*tolQRA-tolB-pal*::*kan*$^{R}$ | This study |
| MG1655 Δ*exbD*::*frt* | This study |
| MG1655 Δ*exbD*::*frt* Δ*tolQRA*::*kan*$^{R}$ | This study |
| MG1655 Δ*tolA*::*frt* | This study |
| W3110 *pal-mCherry* | Petiti et al. (2019) |
| W3110 *pal-mCherry* Δ*tolQR*::*kan*$^{R}$ | Petiti et al. (2019) |

#### Plasmid construction

pBAD33-*exbBD* was cloned using Gibson assembly with the *exbBD* gene sequence inserted into the original multiple-cloning sites on pBAD33 with the following junctions (start/stop underlined):

pBAD33 5′-CGAATTCGAGCTCGGTAGGAGGTACACA**GTGGGTAATAATTT**-3′ **exbBD**<br>
**exbBD** 5′-**CCGCCAAAGCGAAGTAA**CCTCTAGAGTCGACC-3′ pBAD33

pET23/42-*tonB*$^{TM}$*-tolA* was cloned using Gibson assembly with the *tonB* and *tolA* gene fragments inserted into the original multiple-cloning sites on pET23/42 with the following junctions (start/stop underlined):

pET2342 5′- AAGAAGGAGATATACAT**ATGACCCTTGATTTACCTCG**-3′ **tonB**<br>
**tonB**$^{34}$ 5′- **TACCTCGGTACATCAG***GATGAGAATATAGAA*-3′ $^{35}$*tolA*<br>
*tolA* 5′- *CATTGGACTTCAAACCGTAA*CTCGAGCACCACCACCACCA-3′ pET2342

pET23/42-*sfgfp-tonB*$^{TM}$*-tolA* was cloned using Gibson assembly with the *sfgfp* gene fragments inserted into the pET23/42-*tonB-tolA* plasmid with the following junctions (start/stop and linker underlined):

pET2342-tonB-tolA 5′- AAGAAGGAGATATACAT**ATGTCTAAAGGTGAAGAACTGTT**-3′ **sfgfp**<br>
**sfgfp** 5′- **GGATGAGCTCTACAAA**GGATCCACCCTTGATTTACCTCGCCG-3′ pET2342-tonB-tolA

pET23/42-*sfgfp-tolA* was cloned using Gibson assembly with the *sfgfp* and *tolA* gene fragments inserted into the original multiple-cloning sites on pET23/42 with the following junctions (start/stop and linker underlined):

pET2342 5′- AAGAAGGAGATATACAT*ATGTCTAAAGGTGAAGAACTGTT*-3′ *sfgfp*<br>
*sfgfp* 5′- *GGATGAGCTCTACAAA*GGATCC**TCAAAGGCAACCGAACAAAAC**-3′ **tolA**<br>
**tolA** 5′- **CATTGGACTTCAAACCGTAA**CTCGAGCACCACCACCACCA-3′ pET2342

pET23/42-*sfgfp* was cloned using Gibson assembly with the *sfgfp* gene fragment inserted into the original multiple-cloning sites on pET23/42 with the following junctions (start/stop underlined):

pET2342 5′- AAGAAGGAGATATACAT**ATGTCTAAAGGTGAAGAACTGTT**-3′ **sfgfp**<br>
**sfgfp** 5′- **GCATGGATGAGCTCTACAAATAA**CTCGAGCACCACCACCACCA-3′ pET2342

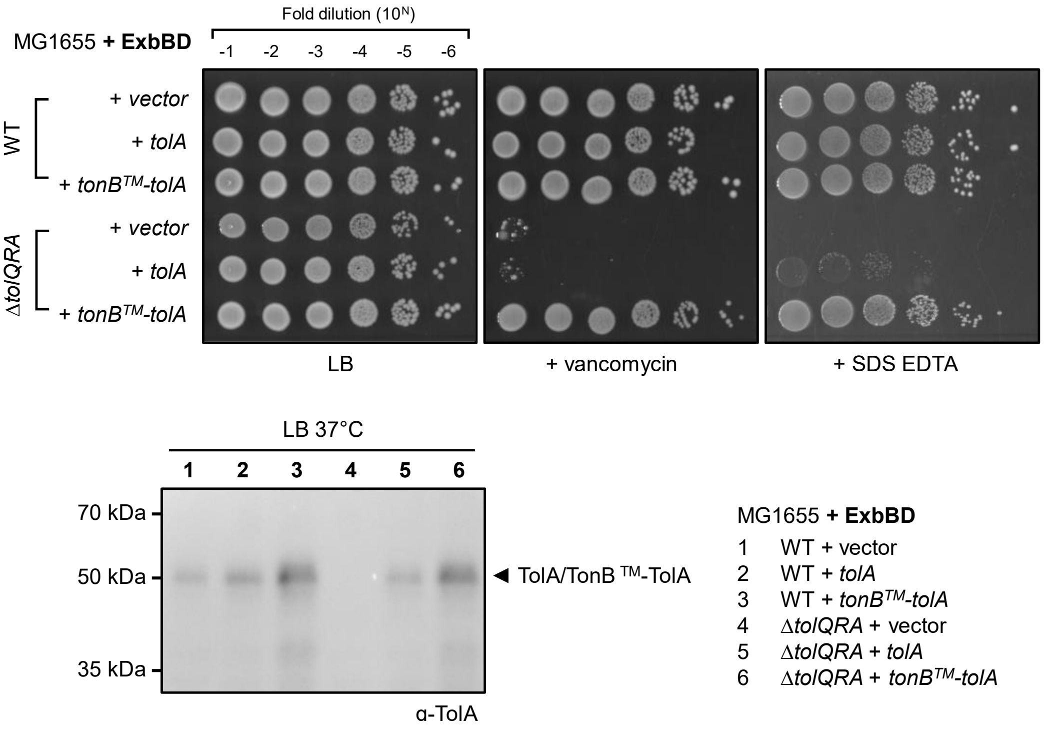


**Figure S1. TonB$^{TM}$-TolA but not wild-type TolA rescue outer membrane defects in strains lacking TolQRA.** (*Top*) Efficiency of plating (EOP) of MG1655 WT and Δ*tolQRA* strains, with either the pET23/42 empty vector or the plasmid expressing full length TolA or TonB$^{TM}$-TolA chimera protein, on LB agar plates supplemented with vancomycin (60 µg/ml) or SDS EDTA (0.3%, 0.3 mM) at 37°C. (*Bottom*) Western blot of the indicated strains grown at 37°C with anti-TolA antibody. TolA and TonB$^{TM}$-TolA are expressed stably from the pET23/42 vector. All strains here express additional copies of ExbBD from a pBAD33 vector.

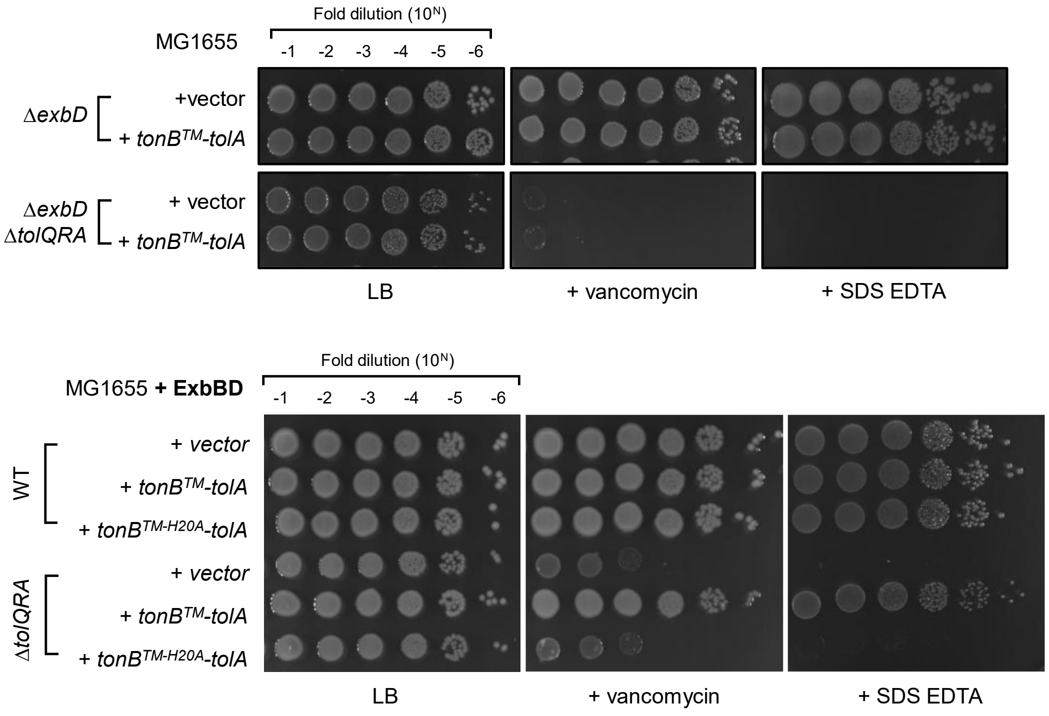


**Figure S2. TonB$^{TM}$-TolA forms a complex with ExbBD for function.** (*Top*) EOP of MG1655 Δ*exbD* and Δ*exbD* Δ*tolQRA* strains, with either the pET23/42 empty vector or the plasmid expressing the TonB$^{TM}$-TolA chimera protein, on LB agar plates supplemented with vancomycin (30 µg/ml) or SDS EDTA (0.3%, 0.3 mM) at 37°C. (*Bottom*) EOP of MG1655 WT and Δ*tolQRA* strains (with additional ExbBD), with either the pET23/42 empty vector or the plasmid expressing TonB$^{TM}$-TolA chimera protein or its H20A variant, on LB agar plates supplemented with vancomycin (60 µg/ml) or SDS EDTA (0.3%, 0.3 mM) at 37°C.

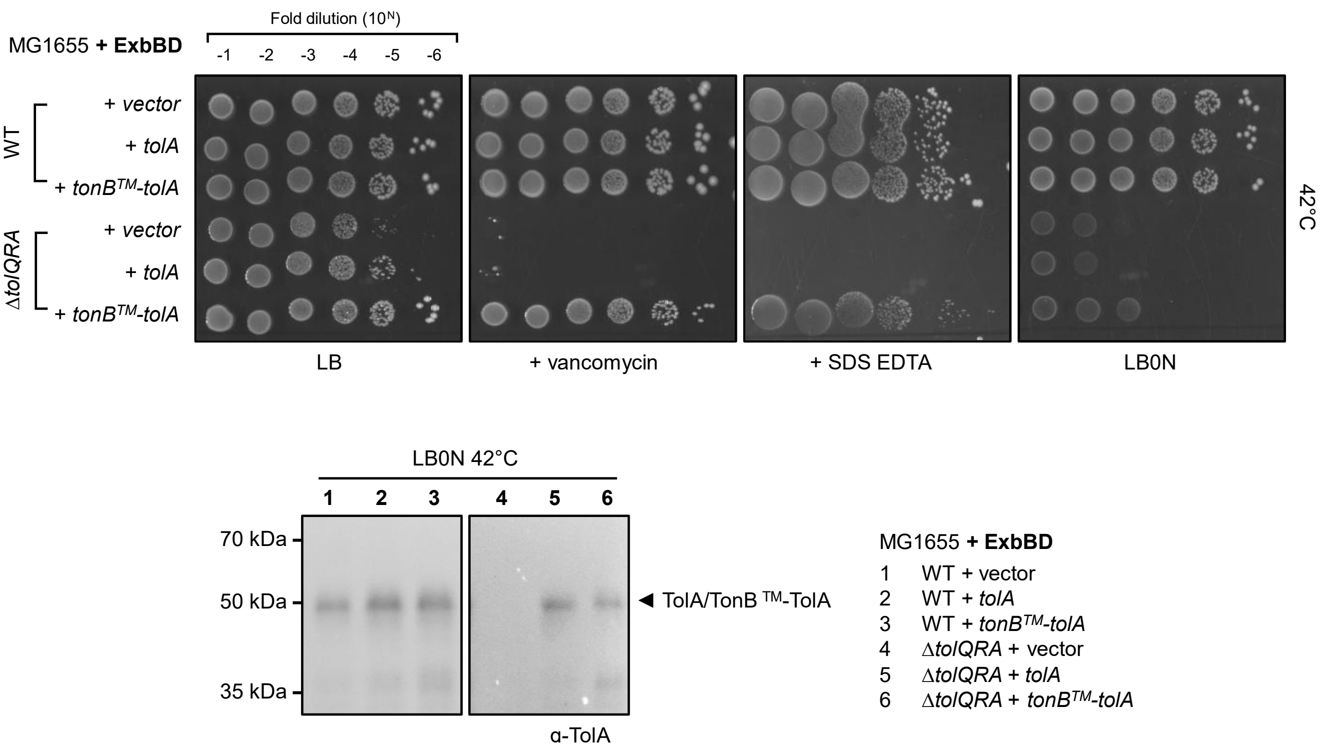


**Figure S3. TonB$^{TM}$-TolA is expressed stably and capable of restoring outer membrane barrier at elevated temperatures.** (*Top*) EOP of MG1655 WT and Δ*tolQRA* strains, with either the pET23/42 empty vector or the plasmid expressing full length TolA or TonB$^{TM}$-TolA chimera protein, on LB agar plates supplemented with vancomycin (30 µg/ml) or SDS EDTA (0.3%, 0.3 mM), or LB0N at 42°C. (*Bottom*) Western blot of the indicated strains grown at 42°C in LB0N with anti-TolA antibody. All strains here express additional copies of ExbBD from a pBAD33 vector.

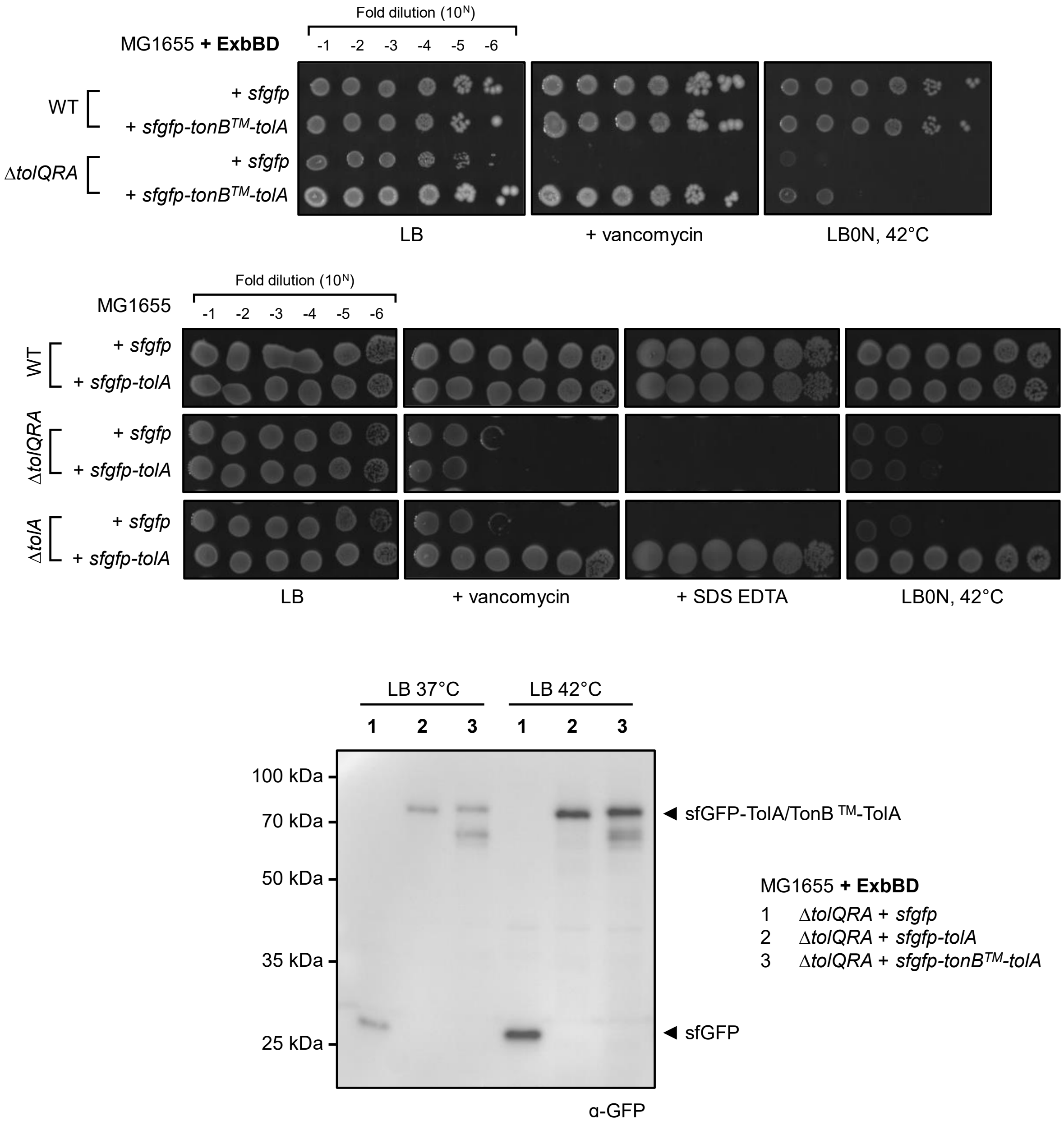


**Figure S4. sfGFP-TonB$^{TM}$-TolA and sfGFP-TolA are functionally identical to their untagged version.** (*Top*) of MG1655 WT and Δ*tolQRA* strains (with additional ExbBD), expressing either sfGFP (negative control) or sfGFP-TonB$^{TM}$-TolA, on LB agar plates supplemented with vancomycin (30 µg/ml) at 37°C, or LB0N at 42°C. (*Middle*) EOP of MG1655 WT, Δ*tolA*, and Δ*tolQRA* strains, expressing either sfGFP or sfGFP-TolA, on LB agar plates supplemented with vancomycin (30 µg/ml), SDS EDTA (0.3%, 0.3 mM) at 37°C or LB0N at 42°C. (*Bottom*) Western blot of the Δ*tolQRA* strains (with additional ExbBD) expressing either sfGFP, sfGFP-TolA, or sfGFP-TonB$^{TM}$-TolA grown in LB at either 37°C or 42°C with anti-GFP antibody.

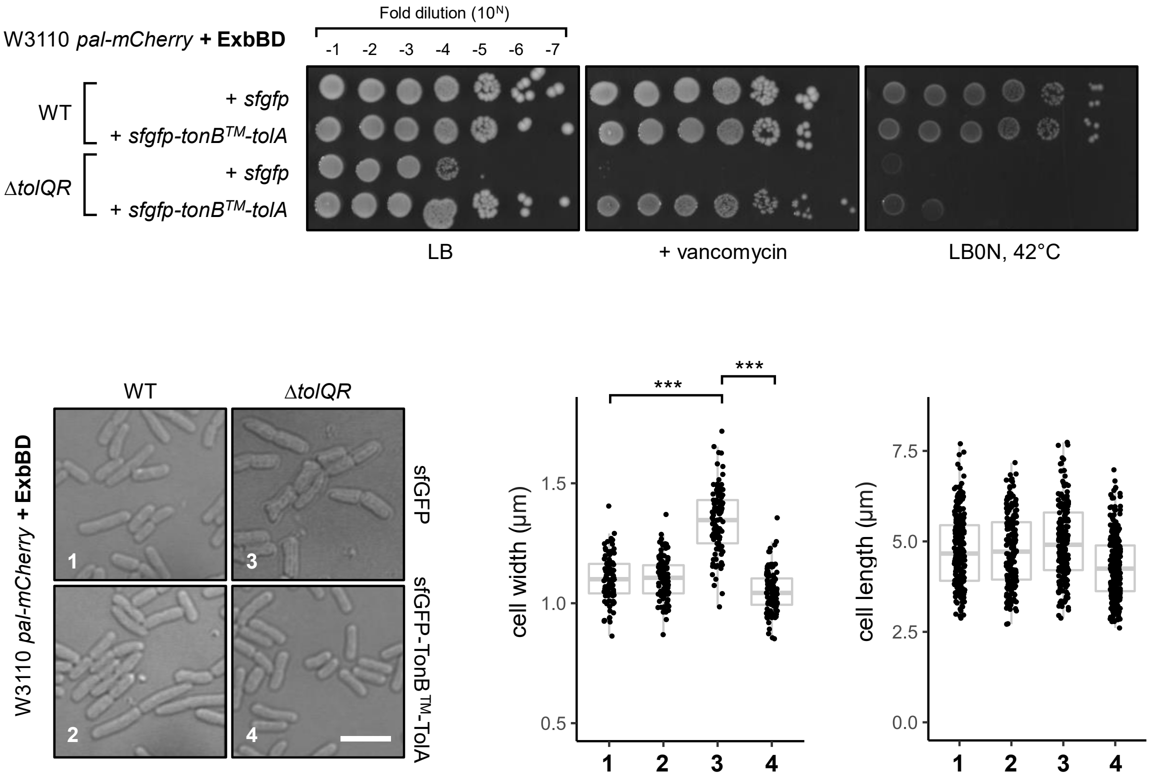


**Figure S5. sfGFP-TonB$^{TM}$-TolA is functional and rescues cell width defects in W3110 *pal-mCherry* strains.** (*Top*) EOP of W3110 *pal-mCherry* WT and Δ*tolQR* strains, expressing either sfGFP or sfGFP-TonB$^{TM}$-TolA, on LB agar plates supplemented with vancomycin (30 µg/ml) at 37°C, or LB0N at 42°C. (*Bottom*) Differential interference contrast (DIC) microscopy images of the same strains shown above. Scale bar represents 5 µm. (*Right*) Quantification of the cell width and length of >100 individual cells of the same strains (numbered) shown in *left* panel. Wilcoxon ranked sum test: ***, $P < 0.0001$. All strains here express additional copies of ExbBD from a pBAD33 vector.
